# Ultrashort-T_2_* mapping at 7 tesla using an optimized pointwise encoding time reduction with radial acquisition (PETRA) sequence at standard and extended echo times

**DOI:** 10.1101/2024.09.05.611365

**Authors:** Carly A. Lockard, Bruce M. Damon, Hacene Serrai

## Abstract

Zero echo time (ZTE) sequences capture signal from tissues with extremely short T_2_* and are useful for qualitative and quantitative imaging of musculoskeletal tissues’ ultrashort-T_2_* components. One such sequence is Pointwise Encoding Time Reduction with Radial Acquisition (PETRA). While this sequence has shown promising results, it has undergone only limited testing at 7 tesla (T). The purpose of this work was to evaluate PETRA at 7T in its standard form and with sequence modifications to allow extended echo times for the purpose of performing ultrashort-T_2_* mapping.

We acquired PETRA images of MnCl_2_ and collagen phantoms and of the knee in eight participants (5 for optimization and 3 for ultrashort-T_2_* mapping assessment; 5 male/3 female, 39 ± 11 years old). Images were acquired using a 1-transmit/28-receive-channel knee coil. Artifacts, signal, signal-to-noise ratio (SNR), ultrashort-T_2_*, the corresponding curve fit quality, and repeatability were assessed. SNR was high in knee tissues at TE = 0.07 msec compared to a conventional-TE sequence (Dual-Echo Steady State with TE = 2.55 msec), with values ranging between 68 to 337 across the assessed tissues for PETRA versus 16 to 30 for the same tissue regions of interest in the conventional-TE series. Acquisition of series for ultrashort-T_2_* maps was feasible at 1.50 mm isotropic acquisition resolution and TE ≤ 0.58 msec. Strong linear correlations were observed between relaxation times and MnCl_2_ concentration, and between signal and collagen concentration. Ultrashort-T_2_* signal decay curve fit R^2^ and repeatability were high for phantom and knee ultrashort-T_2_* <1 msec.

PETRA imaging with minimal artifacts, high SNR, and scan time < 11 minutes was achieved at 7T at high (0.34 mm isotropic) resolution at TE = 0.07 msec and lower resolution (1.52 mm isotropic) at echo times ≤ 0.58 msec. Ultrashort-T_2_* mapping provided acceptable curve-fitting results for substances with sub-millisecond T_2_*.

## Introduction

Conventional magnetic resonance imaging (MRI) sequences provide limited visualization of tissues with short/ultrashort T_2_* values, due to their inability to reduce the echo times (TE) sufficiently to acquire signal from these tissues prior to signal decay. However, ultrashort echo-time (UTE) and zero echo-time (ZTE) MRI sequences are designed to acquire signal from these tissues, enabling qualitative and quantitative evaluations. In the musculoskeletal system, UTE and ZTE imaging are particularly useful for studies of bone, tendon, calcified cartilage, meniscus, and other organs[1–10]. One ZTE sequence that has begun to be applied to these tissues is Pointwise Encoding Time Reduction with Radial Acquisition (PETRA)[3,7,11,12].

The PETRA sequence allows 3D isotropic imaging with TEs under 0.1 msec by applying the readout gradients prior to the radiofrequency pulse and combining radial half-projection filling of the outer portions of k-space with single pointwise Cartesian filling of the central portions of k-space, which are otherwise missed during the transmit/receive switching delay[6,13]. PETRA has been applied at 3T for *in vivo* imaging of the knee with a single TE with a focus on meniscus[11,14], cartilage[14], and the cruciate[14] and collateral ligaments[14]. PETRA has also been applied with dual-echo and inversion-based long-T_2_ suppression strategies[7,13]. Grodzki et al. performed in vivo imaging of the wrist, knee, and foot/ankle using two echo times (TE_1_ ≤ 0.07 msec, TE_2_ = 4.6 msec) at 1.5T to create subtraction images that highlight ultrashort-T_2_* tissues[13], and Li et al. performed in vivo imaging of the lower leg and foot/ankle at 3T with a focus on bone visualization by inversion-based long-T_2_* suppression[7]. PETRA-based spin-lattice in the rotating frame (T_1ρ_) relaxometry has been performed in short-T_2_* tissues of ex vivo bovine joint and tendon specimens and in vivo human knee and ankle tendons and ligaments at 3T[12].

The signal-to-noise ratio (SNR) benefits of UTE and ZTE sequences at 7T compared to lower field strengths have been demonstrated. Cortical bone imaging results with a 3D UTE sequence at 7T demonstrated SNR increases of 1.7 times those achieved at 3T, in spite of reduced T_2_* times at 7T compared to 3T[5]. Human testing of a 3D radial algebraic ZTE sequence on a scanner with dedicated high-performance hardware showed that the high SNR achieved at 7T can be used for high resolution (0.83 mm isotropic) ZTE imaging of the human wrist, knee, and foot/ankle[9]. While other ZTE imaging approaches have applied in vivo in human joints at 7T[9,15], the reported applications of PETRA at 7T in musculoskeletal and other ultrashort-T_2_* tissues have been limited. Other ZTE approaches tested at 7T have utilized algebraic reconstruction to fill the gap left in the center of k-space from the dead-time gap[9,15], while PETRA addresses the gap in the center of k-space using Cartesian single point (pointwise) acquisition, allowing complete filling of the center of k-space with exact values[13]. This allows greater flexibility in the dead-time length/k-space gap, whereas algebraic filling cannot be used if the gap exceeds two or three Nyquist dwells[3,16]. PETRA has been found to achieve lower levels of image artifact compared to other ZTE methods[3]. PETRA has been applied to ex vivo imaging of bovine bone specimens at 7T[3] and is available as a commercial sequence at 7T, but we are unaware of previous literature describing 7T testing and optimization in human joints. Therefore, 7T testing and optimization of the PETRA sequence is needed to evaluate image appearance and potential challenges at this higher field strength.

In addition to minimizing artifacts compared to some other ZTE approaches, the Cartesian single point k-space filling approach that PETRA uses allows image acquisition at different k-space gap sizes, corresponding to a range of TEs, as required to perform ultrashort-T_2_* mapping[3,13]. In addition to single-TE PETRA assessment, it would be beneficial to expand the applications of this sequence to ultrashort-T_2_* mapping and to assess the performance of such mapping at 7T. As mentioned above, PETRA-based T_1ρ_ relaxometry[12] has been performed by adding a T_1ρ_ preparation module and performing multiple scans at a series of spin-lock times. T_1ρ_ relaxometry provides information about proteoglycan content[12]. Similarly, ultrashort-T_2_* relaxometry provides quantitative information about tissue degeneration or other changes, potentially related to disruption of normal collagen organization, in ultrashort-T_2_* tissues[17–19]. Ultrashort-T_2_* mapping has been shown to be particularly sensitive to tissue changes and tissue properties in ultrashort-T_2_* tissues such as deep/calcified cartilage[18,19], meniscus[17,20], tendon[21,22], and ligament[23,24]. However, PETRA-based ultrashort-T_2_* mapping has not yet been assessed.

In this work we present ultrashort-T_2_* mapping results using a modified PETRA sequence which allows an increased TE range of 0.07 to 0.58 msec. Using in vivo knee imaging studies, we performed an initial optimization and addressed the appearance of artifacts at the commercially available sequence settings when using the shortest achievable TE. In addition, we evaluated the feasibility of using the modified extended TE range to perform ultrashort-T_2_* mapping in phantom and in vivo knee images.

## Materials and methods

The product PETRA sequence was modified by extending the maximum allowable TE from 0.10 msec to 1.10 msec. The increase in TE values required an increase in the size of the inner k-space portion filled with single pointwise Cartesian approach and a reduction in the outer k-space filled using radial sampling. Together, these changes increased the signal acquisition time, and therefore also the TE. We limited our maximum time for a single scan to 12 minutes for participant tolerance and determined that 0.58 was the maximum TE feasible within this scan time (details provided in the next section). The modified PETRA sequence was implemented on a 7T MAGNETOM Terra (Siemens, Erlangen, Germany) to collect phantom and knee data using a 1-transmit/28-receive-channel knee coil (Quality Electrodynamics, Mayfield Village, Ohio, USA).

### Multi-echo image optimization for ultrashort-T_2_* mapping

The multi-echo image optimization goals included achieving the necessary levels of SNR and artifact reduction to allow ultrashort-T_2_* mapping within clinically feasible scan times, reported as minutes:seconds, and at the highest practically achievable spatial resolution.

#### In vivo knee imaging

Eight study participants were recruited between November 19, 2022 and June 15, 2024. The study was approved by the institutional review board, and written informed consent was obtained from all participants. For the initial optimization studies, five healthy participants (two male, three female, 42 ± 13 years old) were scanned using the modified PETRA sequence for initial optimization. Dielectric pads (7TNS Neuro Set, Multiwave Imaging, Marseille, France) were wrapped around the anterior and posterior surfaces of the knee. All PETRA scans used a single excitation and a flip angle of 6°, and the system transmit/receive reference amplitude was adjusted to 170 V to avoid excessive specific absorption rate. Initial imaging was performed with spatial resolutions ranging from 0.32 to 1.56 mm isotropic and TR values ranging from 3.99 to 8.28 msec. Data with and without chemical shift selective fat saturation[13] were collected to test the effect of fat suppression on data quality. Optimized parameters were selected for 1) a high-resolution series with a single short TE and for 2) a lower-resolution series that made acquisition at extended TE values up to 0.58 msec feasible, given the allowable scan time threshold of 12 minutes. The scan parameters listed in S1 Table were selected as providing the best balance between desired resolution, contrast, and scan time. For all knees, a conventional-TE DESS sequence (repetition time (TR) = 8.68 msec, TE = 2.55 msec, resolution 0.30 mm isotropic, fat suppressed) was also acquired for comparison.

### Ultrashort-T_2_* mapping

#### MnCl_2_ phantom

A MnCl_2_ phantom was used to perform initial evaluation of the ultrashort-T_2_* mapping protocol using scan parameters based on the initial knee optimization work. The MnCl_2_ phantom contained ten 5 mL centrifuge tubes having 0.02 - 27.07 mM MnCl_2_ tetrahydrate and 30 mM NaCl in deionized water[25]. The tubes were arranged within a cylindrical plastic container sized to fit within the knee coil. Plastic supports were used to keep the tubes stationary in the central volume of the container.

Two scan sessions were performed. In the first session, the phantom was imaged with a conventional-TE DESS sequence (TR = 8.68 msec, TE = 2.55 msec, resolution 0.30 mm isotropic, fat suppressed) and with the PETRA sequence using: matrix = 96; FOV = 150mm isotropic; 29,000 radial views; voxel size = 1.56 mm isotropic; flip angle = 6°; receive bandwidth = 365 Hz/pixel; fat suppressed; TR = 7.13 msec; and TE values of 0.07, 0.09, 0.12, 0.16, 0.20, 0.24, 0.30, 0.40, and 0.50 msec. To test within-day repeatability, the TE = 0.07 and 0.50 msec scans were repeated. Acquisition time ranged from 4-5 minutes for TE = 0.07-0.40 msec and was 7 minutes for TE = 0.50 msec. The second session replicated the first session, apart from repeating the TE = 0.07 and 0.50 msec scans.

#### Collagen phantom

The collagen phantom contained nine 5 mL centrifuge tubes filled with water-collagen solutions having different collagen (chicken hydrolyzed collagen powder, BulkSupplements, Henderson, NV, USA) weight/volume (W/V) percentages (0, 5, 10, 15, 20, 25, 30, 35, and 40%). The tubes were arranged and supported within a cylindrical container in a similar manner to the MnCl_2_ phantom. The container was filled with approximately 1L of 30 mM NaCl in deionized water for coil loading.

Two scan sessions were performed, spaced six weeks apart. During both sessions, PETRA series were acquired using the following parameters: total scan time = 5:21; matrix = 160; FOV = 160 mm isotropic; 80,000 radial views; voxel size = 1.00 mm isotropic; flip angle = 6°; bandwidth = 365 Hz/pixel; TR = 4.00 msec; TE = 0.07 msec. During the first scan session, the phantom was imaged with the phantom in two physical orientations. The phantom was rotated about its long axis (which was aligned with the scanner’s z-axis for both scans) by 180° relative to its initial orientation for the second scan. This was done to test for spatial positioning effects on signal homogeneity. During the first scan session, scans with TE values 0.09, 0.12, 0.16, 0.20, 0.24, 0.30, 0.40, and 0.50 msec were also acquired to allow ultrashort-T_2_* mapping. Conventional T_2_* mapping using an axial plane multi-echo GRE sequence was also performed during the first scan session (matrix = 132×134 (264×268 interpolated); FOV = 157×159 mm; pixel size = 0.595×0.593 mm in-plane; slice thickness = 1.20 mm; flip angle = 15°; bandwidth = 600 Hz/pixel; fat suppressed; TR = 37.00 msec; TE = 1.60, 3.81, 5.82, 7.83, 9.84, 11.85, 23.00 msec).

#### In vivo knee imaging

PETRA imaging, using the parameters listed for the low resolution series in S1 Table and with seven TE values (0.07, 0.09, 0.12, 0.16, 0.22, 0.30, and 0.58 msec; total cumulative scan time for all TE values = 44:03), were performed in three male participants (33 ± 5 years old) with no history of knee injury to test the feasibility of performing ultrashort-T_2_* mapping *in vivo*. Ultrashort-T_2_* mapping repeatability data (two scans separated by 10 days) were acquired for one of these participants.

### Image analysis procedures

#### General

For all images, 3D Slicer[26] was used to perform segmentation and MATLAB (R2024a, The Mathworks, Natick, MA) was used to measure/calculate results within the desired regions of interest (ROIs).

#### Image Segmentation

ROIs in the knee and image background were manually drawn in 3D Slicer by a researcher with 8 years of knee MRI segmentation experience. For all tissues except cartilage, skin, and cortical bone, the ROIs included the entire tissue of interest in at least one slice where the tissue was clearly visualized. The cartilage segmentation excluded the most medial and lateral slices, where partial volume averaging or blurring occurred. The skin and cortical bone segmentations were limited to central regions within a small set of slices, to avoid thinner tissue regions relative to the scan resolution and the resulting partial volume averaging and the blurring seen in the most medial and lateral slices. For all tissues, regions near the proximal and distal borders of the image were excluded, to avoid signal inhomogeneity near the edges of the coil’s sensitive volume.

For the phantoms, segmentation was performed using the TE = 0.07 msec images. Circular ROIs of 7 mm diameter, centered within each 15 mm diameter centrifuge tube and avoiding regions with air bubbles, were drawn in on six slices within the straight section of the tube.

#### Analysis of Signal Behavior

In the knee of one participant, the SNR was measured in the cortical bone, patellar tendon, posterior cruciate ligament, anterior cruciate ligament, cartilage, and skin in the fat-suppressed low resolution (1.52 mm isotropic) and high resolution (0.34 mm isotropic), TE = 0.07 msec PETRA series. The mean signal in the ROIs described above was calculated. PETRA image noise was measured by calculating the difference between two subsequently acquired identical series, and then calculating the standard deviation of the difference image divided by √2 to correct for the difference operation[27].

A custom MATLAB script was used to analyze the signal-concentration relationship at TE = 0.07 msec (collagen phantom) and to calculate the ultrashort-T_2_* maps using the TE = 0.07-0.50 msec data (MnCl_2_ phantom and knee images).

#### Ultrashort T2* Fitting

For the MnCl_2_ phantom studies, the ultrashort-T_2_* maps were calculated from 1.50 mm isotropic, fat-suppressed PETRA at nine different TE values (0.07-0.50 msec) using voxel-wise monoexponential fitting. A noise term was not included because of the high SNR. The coefficient of determination, R^2^, for the ultrashort-T_2_* fitting was calculated for each voxel. The median and interquartile range of the ultrashort-T_2_* for the MnCl_2_ phantom and signal mean ± standard deviation for the collagen phantom were calculated. The relationships between ultrashort-T_2_* and MnCl_2_ concentration and between signal intensity at TE = 0.07 msec and collagen concentration were evaluated.

For in vivo studies, ultrashort-T_2_* values were calculated using the same process described for the MnCl_2_ phantom, except that the 1.52 mm isotropic, fat-suppressed PETRA at seven TE values (0.07-0.58 msec) were used (due to the need for a shorter total scan time for in vivo imaging). Similar fitting with a subset of three TE values (0.07, 0.12, and 0.30 msec) was performed with the goal of enabling a reduced total scan time (16:43), thereby reducing the chances of participant motion between series and increasing the practical feasibility. In addition, in-plane frequency-domain interpolation was performed in MATLAB prior to segmentation and ultrashort-T_2_* fitting to allow more precise delineation of the knee structures (resulting in a sagittal-plane series interpolated resolution of 0.76 x 0.76 x 1.52 mm). Ultrashort-T_2_* values for which R^2^ was < 0.5 were excluded. The ultrashort-T_2_* values and R^2^ values were measured in the cortical bone, patellar tendon, meniscus, posterior cruciate ligament, anterior cruciate ligament, cartilage, and skin. *X*^2^ was also calculated to allow more robust comparison between goodness-of-fit between the fitting using seven TE values and three TE values.

## Results

### Multi-echo image optimization for ultrashort-T_2_* mapping

#### In vivo knee imaging

Fig 1 shows an example high-resolution PETRA knee image, with comparison to a dual-echo steady state (DESS) conventional TE sequence image. The PETRA images were obtained with high SNR, even for ultrashort-T_2_* tissues. As listed in Table 1, the SNR in the cortical bone, patellar tendon, meniscus, posterior cruciate ligament, anterior cruciate ligament, cartilage, and skin in the TE = 0.07 msec, 1.52 mm isotropic resolution, TR = 4.21 msec series ranged from 68 to 337 (total scan time, 5:20). In the 0.34 isotropic resolution, TR = 7.07 msec series (Fig 1), the same SNR measurements ranged from 23 to 75 (total scan time, 6:55). Note that the PETRA series at the two resolutions were acquired with different TR, so the SNR does not follow the usual relationship with voxel size between the two series. For comparison, the SNRs in these regions for the 0.30 mm isotropic resolution conventional-TE DESS series ranged from 16 to 30.

**Fig 1.**
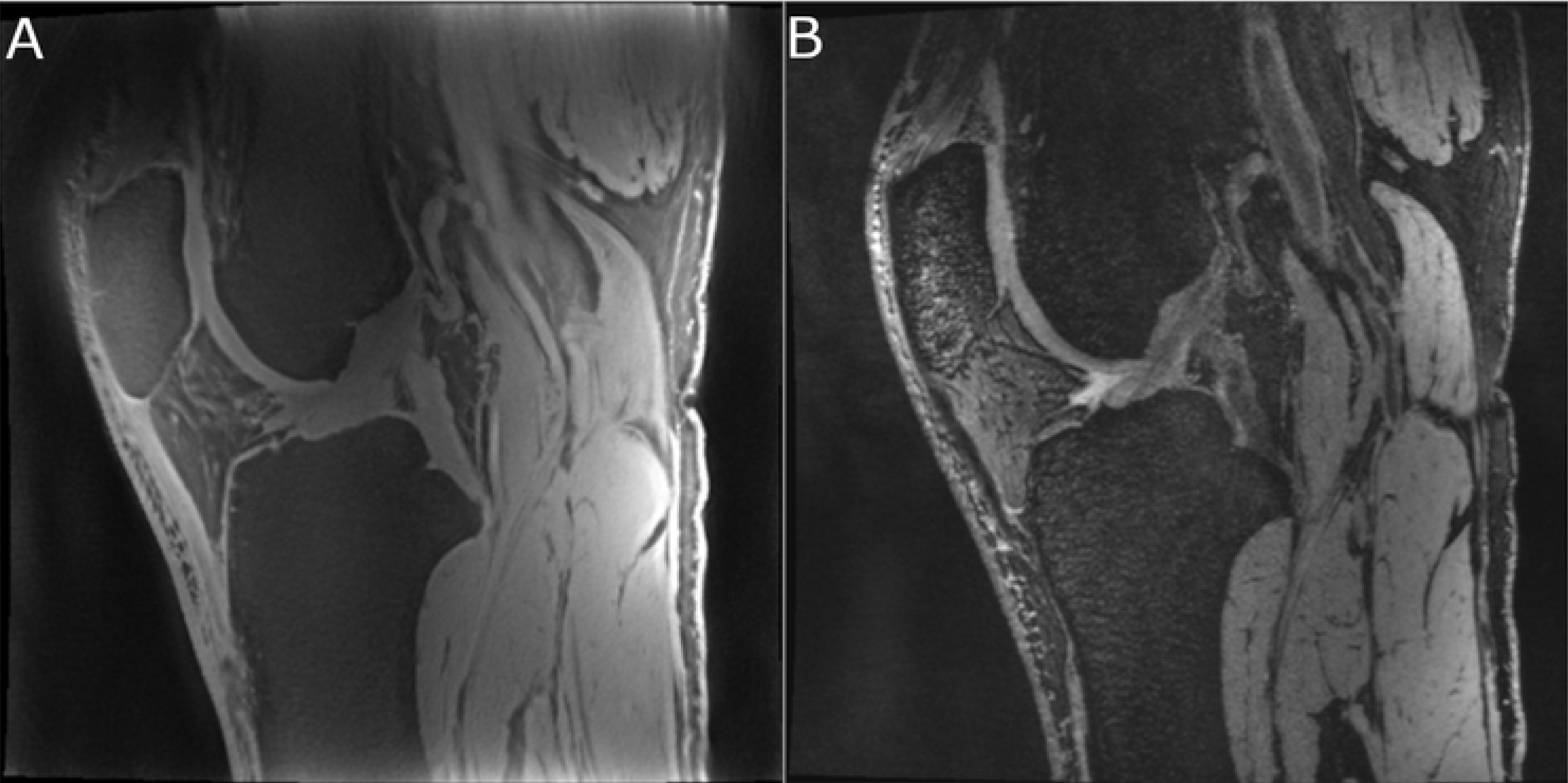
Example high-resolution PETRA knee image, with comparison to a conventional-TE DESS sequence image. Sagittal images of one participant’s knee from the fat-suppressed PETRA series at 0.34 mm isotropic resolution (A) and from the fat-suppressed dual-echo steady state (DESS) at 0.30 mm isotropic resolution (B). These images illustrate the difference in signal, and resulting differences in contrast between conventional and zero echo-time sequences, demonstrated in the patellar tendon, cruciate ligaments, and other ultrashort-T_2_* tissues in these images.

**Table 1.** SNR measurements for PETRA at TE = 0.07 msec for knee tissue ROIs at two resolutions.

|  | SNR at TE = 0.07<br>msec/TR = 7.07 msec<br>for 0.34 mm isotropic<br>resolution | SNR at TE = 0.07<br>msec/TR = 4.21 msec<br>for 1.52 mm isotropic<br>resolution |
| --- | --- | --- |
| Knee tissue |  |  |
| Cortical bone | 23 | 68 |
| Patellar tendon | 58 | 183 |
| Meniscus | 36 | 139 |
| Posterior cruciate ligament | 39 | 163 |
| Anterior cruciate ligament | 34 | 143 |
| Cartilage | 52 | 172 |
| Skin | 75 | 337 |

Comparing the images acquired with and without fat suppression revealed that fat suppression reduced the appearance of off-resonance artifacts and improved the visualization of short-T_2_* tissues (Fig 2). Images with 0.34 mm isotropic resolution were acquired in scan times of 6:55 when acquired with fat suppression and 6:00 when acquired without fat suppression. Figure 2 shows example low-resolution images acquired at TE = 0.07 and 0.50 msec and high-resolution images acquired at TE = 0.07 msec, with and without fat suppression.

**Fig 2.**
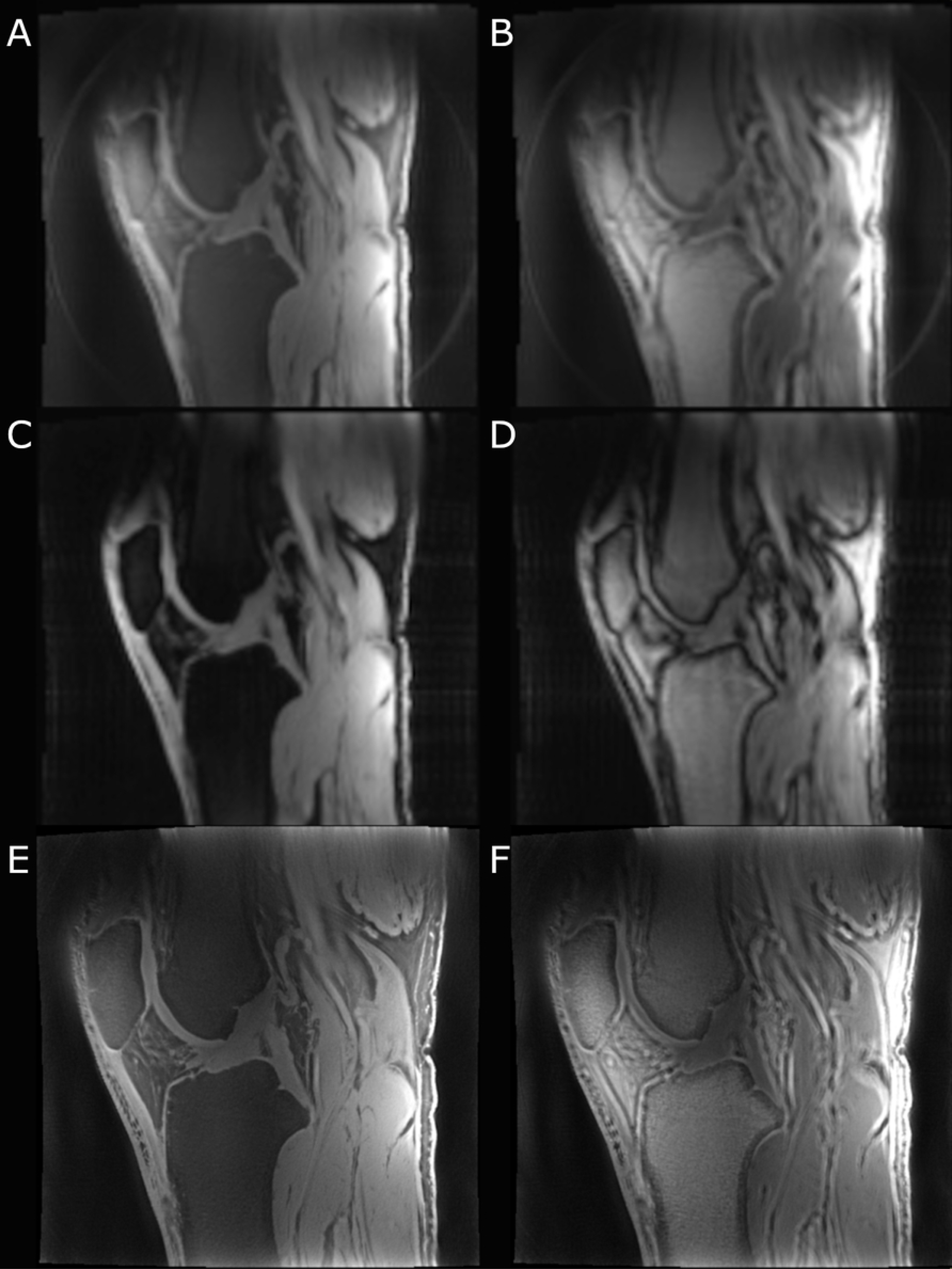
Sagittal PETRA images all acquired in a single participant knee, acquired using the parameters listed in Supplementary Table 1. Low-resolution images acquired at TE = 0.07 msec with (A) and without (B) fat suppression, at TE = 0.58 msec with (C) and without (D) fat suppression, and high-resolution images at TE = 0.07 msec with (E) and without (F) fat suppression. Note the off-resonance artifacts appearing as double lines and thick low/high signal borders around tissue interfaces (e.g. patellar tendon-infrapatellar fat pad interface, muscle-fat interfaces, etc.).

To achieve acquisitions at longer TE values without exceeding 12 minutes per scan, the resolution needed to be set to 1.52 mm isotropic. At this resolution with 70,000 radial views and TR = 4.21 msec the scan time was under 12 minutes at TE values up to 0.58 msec (scan time = 10:55, with fat suppression).

### Ultrashort-T_2_* mapping

#### MnCl_2_ phantom

The ultrashort-T_2_* measurements, ultrashort-T_2_* curve fit *R^2^* values, and interscan coefficients of variation (CV) for each phantom solution ROI for two sessions are shown in Table 2. Example ultrashort-T_2_* curve fits are shown in Fig 3A. There was a significant linear correlation between 1/T_2_*_ultrashort_ values and MnCl_2_ concentration (*r* = 0.99 and *p* <0.0001 for both scans), as shown in Fig 3B. Mean *R^2^* dropped sharply for MnCl_2_ concentrations below 3.6 mM, and at the lowest concentration (0.02 mM) a large number of very high or even negative T_2_* values was calculated due to the lack of signal decay measured over the range of acquired TEs, leading to poor fit quality and the very large interquartile range for that measurement. The scan 1 and scan 2 slopes and intercepts were calculated using just the results for the solution concentrations with average *R^2^* for ultrashort-T_2_* fit ≥ 0.90 (which corresponded to 5.4 - 27.07 mM, with ultrashort-T_2_* values of ≤ 1.1 msec). The scan 1 and scan 2 slopes were 141.77 and 135.81 seconds^-1^/mM (8% CV between scans), and the intercepts were 211.16 and 218.03 seconds^-1^ (9% CV between scans).

**Fig 3.**
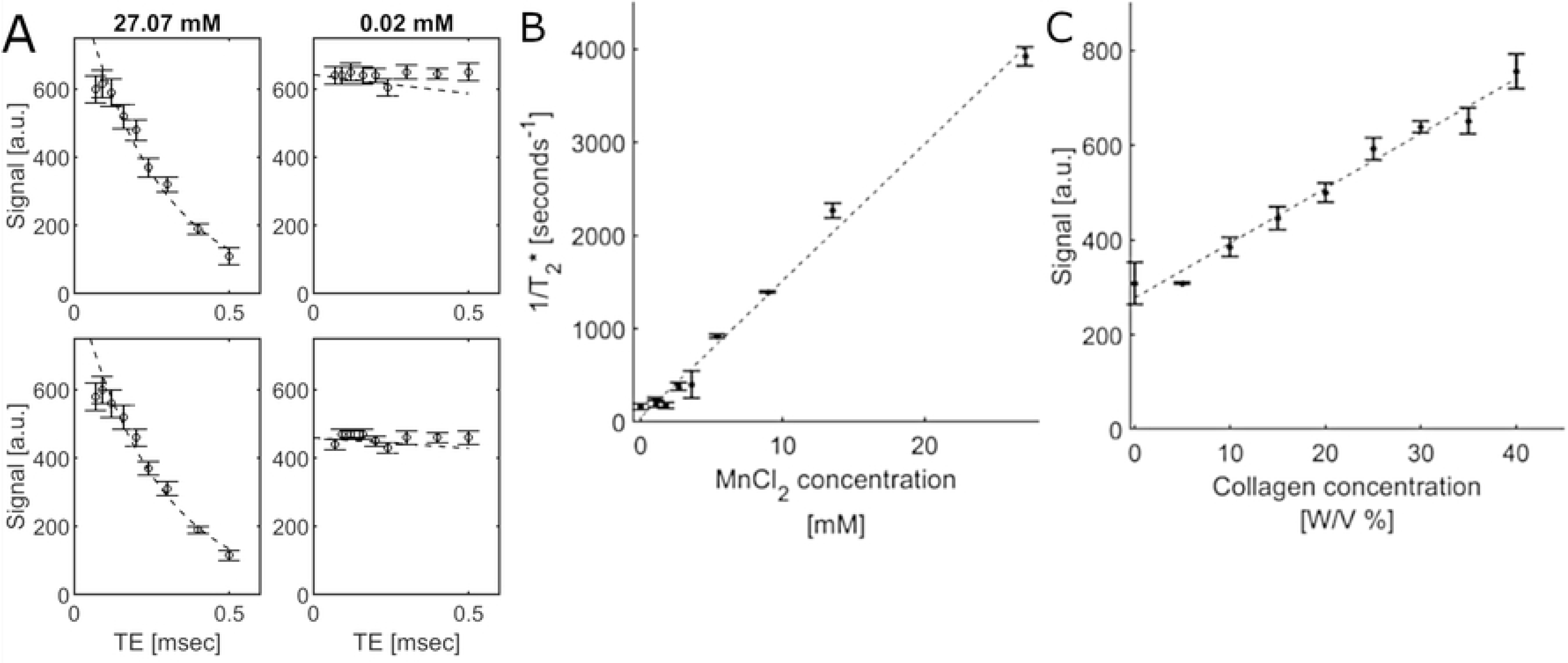
MnCl_2_ phantom plots. Plots showing examples of median measured (round markers) and fitted signal (dashed line) based on estimated ultrashort-T_2_* signal versus TE for three MnCl_2_ phantom solutions from two (top and bottom plots for each region) scans (a), the relationship between mean 1/T_2_* and MnCl_2_ concentration, averaged over two scans (b), the relationship between mean signal and collagen concentration at TE = 0.07 msec, averaged over three scans (c). Error bars represent the interquartile range within each subregion for signal versus TE plots and the standard deviation between scans for the 1/T_2_* and signal versus concentration plots.

**Table 2.** Calculated ultrashort-T_2_* values and between-scan CV, and ultrashort-T_2_* fit R^2^ from two scans for the MnCl_2_ phantom.

| Phantom $\text{MnCl}_2$<br>solution<br>concentration [mM] | Scan 1 $T_2^*$ median<br>(interquartile<br>range) [msec] | Scan 2 $T_2^*$ median<br>(interquartile<br>range) [msec] | Between-scan $T_2^*$<br>CV | Scan 1<br>mean $R^2$ | Scan 2<br>mean $R^2$ |
| --- | --- | --- | --- | --- | --- |
| 27.07 | 0.3 (0.1) | 0.3 (0.1) | 2% | 0.96 | 0.97 |
| 13.53 | 0.4 (0.0) | 0.5 (0.1) | 3% | 0.98 | 0.97 |
| 9.01 | 0.7 (0.1) | 0.7 (0.1) | 0% | 0.90 | 0.91 |
| 5.4 | 1.0 (0.2) | 1.1 (0.2) | 2% | 0.97 | 0.95 |
| 3.6 | 3.4 (3.7) | 2.0 (0.6) | 26% | 0.51 | 0.85 |
| 2.7 | 2.9 (2.7) | 2.5 (0.6) | 8% | 0.54 | 0.74 |
| 1.79 | 5.0 (2.2) | 6.5 (1.9) | 13% | 0.35 | 0.24 |
| 1.12 | 4.7 (0.7) | 6.1 (2.2) | 13% | 0.38 | 0.30 |
| 1.07 | 4.0 (2.2) | 4.4 (2.0) | 5% | 0.56 | 0.64 |
| 0.02 | 5.6 (37.6) | 7.3 (25.3) | 13% | 0.13 | 0.14 |

#### Collagen phantom

For the collagen phantom, the mean conventional-T_2_* values for all W/V percentages greatly exceeded the ultrashort-T_2_* upper limit, being 23 msec for 40% collagen to 330 msec for 5% collagen; therefore ultrashort-T_2_* mapping was not evaluated. There was a significant linear correlation between collagen concentration and signal intensity in the TE = 0.07 msec images (*r* = 0.98 on average, *p* <0.0001). The individual signal measurements, linear fitting parameters, and correlation statistics are shown in Table 3. Fig 3C shows this relationship for the average of the three TE = 0.07 msec PETRA series. The between-scan and between-orientation signal and linear fitting parameter CV were all ≥ 0.05 except for 0% collagen signal (Table 3).

**Table 3.** Signal measurements for each collagen concentration from three scans of the collagen phantom and the resulting signal versus concentration linear fitting parameters and correlation statistics.

|  | Scan 1, orientation 1 | Scan 1, orientation 2 | Scan 2, orientation 1 | CV |
| --- | --- | --- | --- | --- |
| Collagen W/V<br>percentage [%] | Mean signal (standard deviation) measurements [a.u.] |  |  |  |
| 0 | 301 (32) | 268 (16) | 355 (20) | 14% |
| 5 | 307 (16) | 309 (15) | 310 (10) | 1% |
| 10 | 389 (13) | 403 (23) | 362 (10) | 5% |
| 15 | 463 (11) | 455 (25) | 417 (10) | 5% |
| 20 | 520 (21) | 481 (25) | 496 (13) | 4% |
| 25 | 604 (19) | 564 (19) | 607 (24) | 4% |
| 30 | 637 (16) | 627 (19) | 651 (23) | 2% |
| 35 | 680 (21) | 646 (14) | 625 (9) | 4% |
| 40 | 792 (30) | 719 (43) | 752 (13) | 5% |
| Linear fit and correlation parameter values |  |  |  |  |
| Slope [a.u.%] | 12.40 | 11.26 | 11.01 | 6% |
| Intercept [a.u.] | 273.34 | 271.74 | 288.32 | 3% |
| r | 0.99 | 0.99 | 0.96 |  |
| p | <0.0001 | <0.0001 | <0.0001 |  |

#### In vivo knee imaging

Acquiring images with different TEs but other parameters held constant allowed creation of ultrashort-T_2_* maps. Fig 4 shows an ultrashort-T_2_* map and corresponding *R^2^* map for one knee for two different scan sessions. Ultrashort-T_2_* measurements and fitting information for the cortical bone, patellar tendon, meniscus, posterior cruciate ligament, anterior cruciate ligament, cartilage, and skin are shown in Table 4. Scan-rescan results for one participant and interscan coefficients of variation are shown in Table 5. Results obtained using all seven acquired TE values and using a subset of three TE values are presented for both tables. Example T_2_* curves are shown in Fig 5.

**Fig 4.**
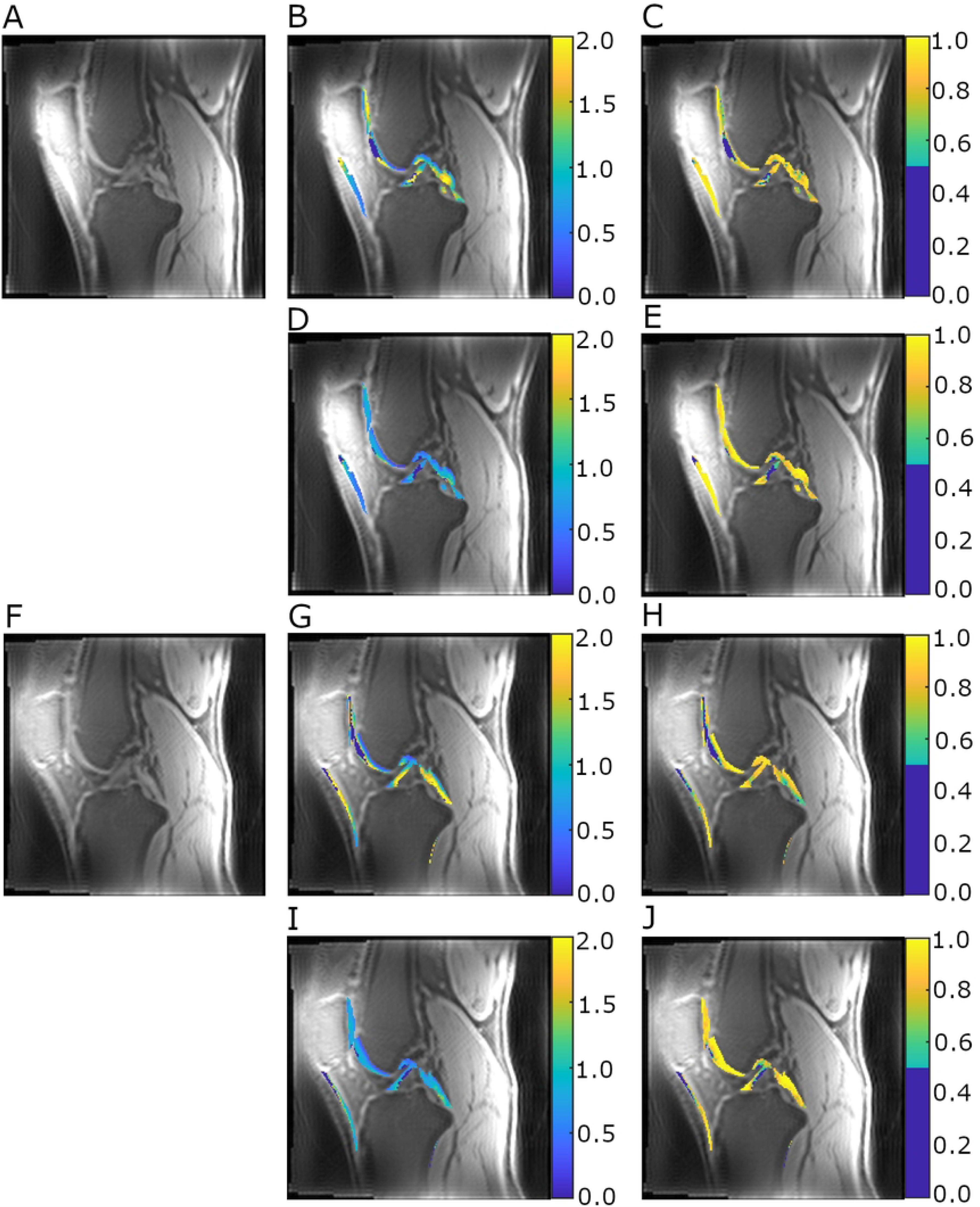
Ultrashort-T_2_* map and corresponding R^2^ map for one knee for two different scan sessions. Example PETRA images at TE = 0.07 msec for one participant’s knee scanned in two sessions (A, F), corresponding calculated ultrashort-T_2_* maps in the masked ROIs in msec (B and D for scan 1 using seven and three TE values, respectively; G and I for scan 2 using seven and three TE values, respectively) and *R^2^* maps in the masked ROIs (C and E for scan 1 using seven and three TE values, respectively; H and J for scan 2 using seven and three TE values, respectively). The *R^2^* value threshold was set to 0.5 in all cases, with voxel ultrashort-T_2_* values set to 0 when *R^2^* for the voxel was below this threshold.

**Fig 5.**
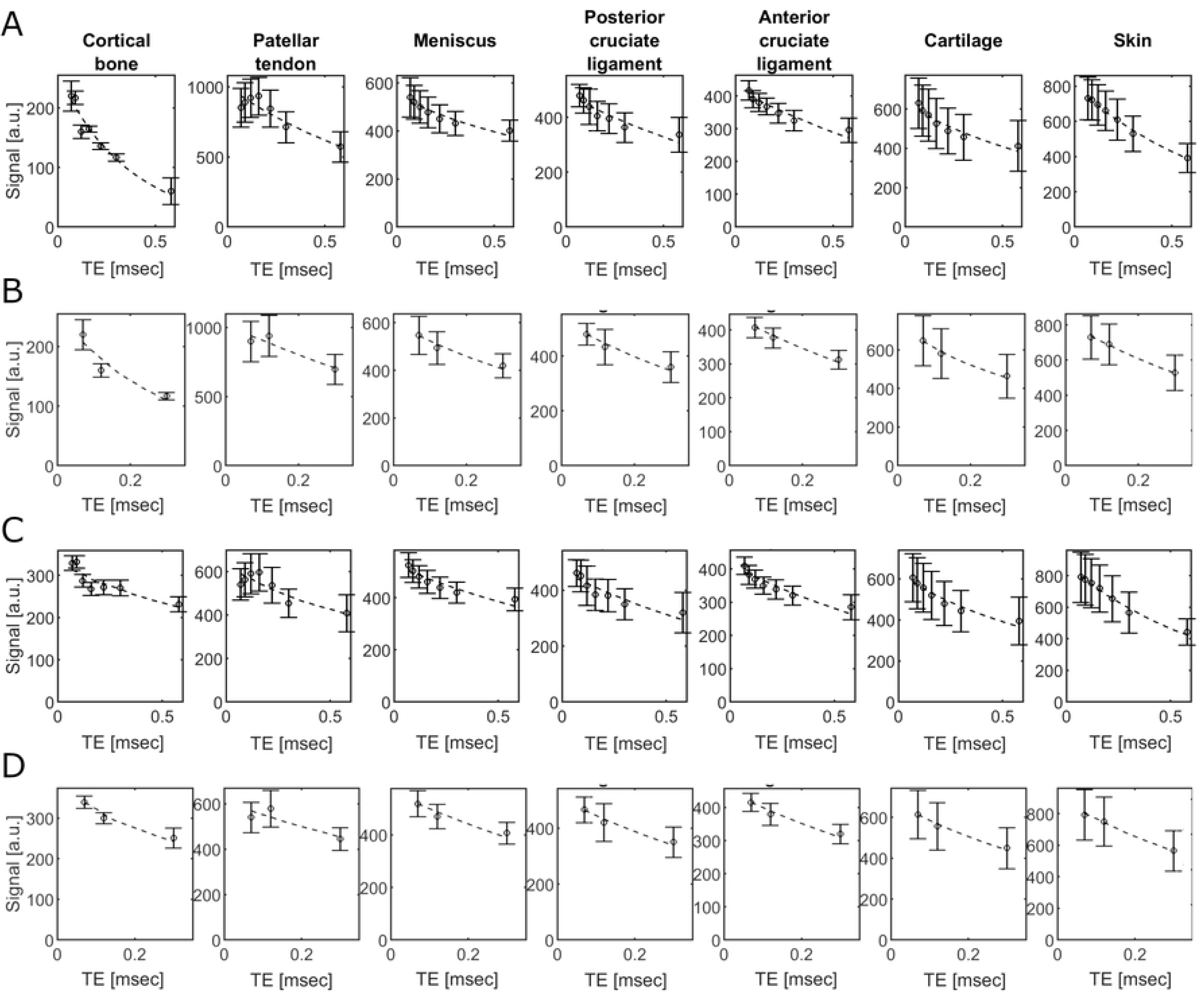
Example T_2_* curves. Plots of the median measured (round markers) and fitted (dashed line) signal based on estimated ultrashort-T_2_* signal versus TE for the measured knee tissue regions from two scans of a single participant’s knee (10 days between scans). The plots for scan 1 are shown in rows A (signal decay curves fitted to all seven TE values) and B (signal decay curves fitted to a subset of three TE values) and the same plots for scan 2 are shown in rows C and D. Error bars represent the interquartile range within each subregion.

**Table 4.** Knee tissue calculated median and interquartile range (IQR) ultrashort-T_2_* values, ultrashort-T_2_* fit *R*^2^, and ultrashort-T_2_* fit *X*^2^ for single scans for three participants. The results calculated using all seven TE values and the subset of three TE values are presented.

| Knee tissue | Median ultrashort-$T_2^*$ (IQR) [msec] | Range of percent voxels excluded for $R^2 < 0.5$ [%] | Median $R^2$ (IQR) | Median $X^2$ (IQR) |
| --- | --- | --- | --- | --- |
| Results when fitting to seven TE values |  |  |  |  |
| Cortical bone | 0.7 (0.5) | 0 - 30 | 0.93 (0.05) | 6.36 (2.83) |
| Patellar tendon | 1.1 (0.1) | 7 - 25 | 0.92 (0.07) | 5.78 (7.49) |
| Meniscus | 1.8 (0.3) | 37 - 42 | 0.77 (0.03) | 8.41 (0.32) |
| Posterior cruciate ligament | 1.3 (0.5) | 6 - 24 | 0.82 (0.01) | 8.79 (3.42) |
| Anterior cruciate ligament | 1.3 (0.5) | 29 - 60 | 0.81 (0.06) | 9.89 (2.74) |
| Cartilage | 1.0 (0.1) | 10 - 25 | 0.91 (0.05) | 10.54 (5.93) |
| Skin | 0.8 (0.1) | 0 - 3 | 0.98 (0.01) | 3.73 (1.14) |
| Results when fitting to three TE values |  |  |  |  |
| Cortical bone | 0.7 (0.2) | 0 - 17 | 0.87 (0.01) | 1.20 (1.28) |
| Patellar tendon | 0.8 (0.1) | 26 - 49 | 0.97 (0.06) | 1.03 (2.30) |
| Meniscus | 0.9 (0.1) | 9 - 27 | 0.88 (0.05) | 1.98 (0.43) |
| Posterior cruciate ligament | 0.8 (0.1) | 3 - 51 | 0.92 (0.09) | 1.59 (0.40) |
| Anterior cruciate ligament | 0.7 (0.1) | 16 - 47 | 0.92 (0.04) | 1.62 (1.66) |
| Cartilage | 0.7 (0.0) | 6 - 11 | 0.96 (0.02) | 0.97 (0.70) |
| Skin | 0.7 (0.1) | 0 - 1 | 0.98 (0.01) | 0.42 (0.09) |

**Table 5.** Scan 1 and scan 2 single-participant ROI median and interquartile range (IQR) ultrashort-T_2_* values and between-scan coefficient of variation (CV) calculated from the two scan sessions for one participant. The results calculated using all seven TE values and the subset of three TE values are presented.

| Knee tissue | Scan 1 median<br>ultrashort-T<sub>2</sub>*<br>(IQR) within each<br>ROI [msec] | Scan 2 median<br>ultrashort-T<sub>2</sub>*<br>(IQR) within each<br>ROI [msec] | Scan-rescan ultrashort-T<sub>2</sub>*<br>CV |
| --- | --- | --- | --- |
| Results when fitting to seven TE values |  |  |  |
| Cortical bone | 0.4 (0.1) | 1.7 (1.2) | 93% |
| Patellar tendon | 1.3 (0.8) | 1.5 (1.1) | 9% |
| Meniscus | 1.6 (1.0) | 1.7 (0.9) | 7% |
| Posterior cruciate<br>ligament | 1.3 (0.8) | 1.3 (0.8) | 0% |
| Anterior cruciate<br>ligament | 1.3 (0.9) | 1.3 (0.7) | 3% |
| Cartilage | 1.1 (1.1) | 1.2 (0.9) | 6% |
| Skin | 0.8 (0.3) | 0.9 (0.4) | 13% |
| Results when fitting to three TE values |  |  |  |
| Cortical bone | 0.4 (0.1) | 0.7 (0.4) | 50% |
| Patellar tendon | 1.0 (0.5) | 1.1 (0.5) | 9% |
| Meniscus | 0.8 (0.3) | 0.9 (0.3) | 6% |
| Posterior cruciate<br>ligament | 0.7 (0.3) | 0.7 (0.3) | 2% |
| Anterior cruciate<br>ligament | 0.7 (0.3) | 0.8 (0.2) | 4% |
| Cartilage | 0.7 (0.4) | 0.7 (0.3) | 6% |
| Skin | 0.7 (0.2) | 0.7 (0.2) | 2% |

## Discussion

We modified a commercially available ZTE sequence, PETRA[13], to extend the maximum technically allowable echo time to up to 1.10 msec and demonstrate the feasibility of extended TE imaging (up to 0.58 msec) with PETRA at 7T in MnCl_2_ and collagen phantoms and in vivo. PETRA imaging was feasible in vivo within a clinically realistic acquisition times at high resolution (0.34 mm isotropic) at short TE values and lower resolution (1.52 mm isotropic) at the high TE values. In addition to distinguishing between tissues and tissue components with comparatively short T_2_* values from those with long T_2_* values, these modifications further allow advanced applications such as ultrashort-T_2_* mapping from 7T ZTE data.

### Image quality indicators

Higher SNR was measured in all acquired PETRA images than in a conventional TE series at similar resolution. Fat suppression was useful for reducing off-resonance artifact. As previously described[9], coil artifact was observed in a few instances. However, it was outside of the knee volume and therefore did not impact image quality. Furthermore, near-isocenter positioning of the knee allows for minimization of lateral image blurring, as reported previously[28].

### Ultrashort T_2_* mapping

The extension of the maximum TE allows ultrashort-T_2_* mapping by multiple acquisitions for substances with sub-millisecond T_2_* values, demonstrated here using the MnCl_2_ phantom and in vivo knees. The ultrashort-T_2_* fitting results, as quantified by *R^2^*, were excellent for ultrashort-T_2_* ≤ 1.1 msec. For MnCl_2_ samples, there was a strong significant correlation between 1/ultrashort-T_2_* values and concentration. Although this finding is not surprising, given the predictions of fast exchange theory, the linearity of the relaxivity plot provides an important quality verification on the T_2_* mapping capabilities of the sequence.

We simulated collagen-rich tissues using a collagen phantom. Although the collagen concentrations were similar to the range found in typical short-T_2_* tissues, the lack of collagen organization resulted in measured conventional T_2_* values that were longer than in the tissues of interest[29]; therefore, ultrashort-T_2_* mapping was not evaluated. However, we observed a strong correlation between collagen concentration and signal at TE = 0.07 msec, likely due to T_1_ differences between the collagen solutions. Previously reported collagen solution T_1_ values suggest that the expected T_1_ range for our 5%-40% collagen solutions would be around 1.6 – 2.6 sec at 3T[30] and somewhat longer at 7T[31–33]. The T_1_ values in short-T_2_* musculoskeletal and skin tissues with would be expected to be around 330-350 (bound water)/390-1000 (pore water) msec in cortical bone at 4.7 T[34,35], 620-700 msec in tendon at 3T[2,36,37], 819-870 msec in ligament at 3T[37], 870 msec in meniscus at 3T[37], and 1.13 sec in cartilage at 3T[37], and 820-1060 msec in dermis and epidermis of the skin at 1.5T [38], with relatively longer T_1_ values expected for each tissue at 7T. This range of T_1_ values would also be expected to create collagen-dependent signal variations in similarly configured PETRA images. While variations in other structural and compositional tissue properties would prevent across-tissue interpretation of the signal variations as reflecting collagen content, it is possible that within-tissue signal variations could be interpretable as variations in collagen content and could be associated with tissue pathology.

The median ultrashort-T_2_* values were generally similar between the three participants. When comparing ultrashort-T_2_* values calculated using seven versus three TE values we found that the median ultrashort-T_2_* values were lower when only three TE values were used than when seven were used (maximum TE = 0.30 msec versus 0.58 msec). This may reflect higher contributions of shorter T_2_* components when shorter TE values are used, with longer components contributing more to the overall measurement as a greater proportion of longer TE values are included.

For five of the seven tissues assessed, fewer voxels were excluded for poor fit when three TE values were used than when seven TE were used. The mean *X*^2^ values were lower when three TE values were used and between-scan CV was lower for all tissues except the cruciate ligaments, for which the between-scan CV was slightly higher when three TE were used but still <4%. For all tissues except cortical bone, the between-scan CV values were <14% when seven TE values were used and <10% when three TE values were used. The poor between-scan reproducibility of the cortical bone ultrashort-T_2_* mapping may be in part due to the thin ROI relative to the low scan resolution and potential for chemical shift artifact around the bone, and the relatively distal location of the bone ROI relative to the image volume center. Midshaft bone measurements, where the cortical bone is thicker and the bone ROI can be centered in the imaging volume, may be more reproducible. The shorter scan time, better decay curve fit, and higher between-scan reproducibility makes the three TE approach preferable for mapping of the ultrashort-T_2_* components. The addition of a conventional-TE T_2_* sequence would be necessary to capture the long-T_2_* components, when desired, regardless of whether three or seven ultrashort-TE values are used since the maximum TE value is < 1 msec in either case.

Ultrashort-T_2_* *R^2^* values were high in the tendon, cartilage, skin, and cortical bone. For patellar tendon, the median value of 1.1 msec when seven TE were used was similar to the value of 1.57 ± 0.11 msec reported for the short T_2_* component in one patellar tendon sample at 7T[39]. In that study human patellar tendon samples were imaged with their long axes oriented at a range of angles relative to the main magnetic field, illustrating the impact of magic angle effect on tendon monoexponential and bicomponent T_2_* [39]. To minimize differences due to this magic angle effect we compared our results, which were measured at a mean patellar tendon angle of 24° ± 3° relative to the main magnetic field, to the 20° angle results from the Hager study[39]. When three TE were used, our median patellar tendon ultrashort-T_2_* value (0.8 msec) was lower than that previously reported value for the single patellar tendon sample imaged at 20°, but within the range reported for a set of four patellar tendon samples imaged at angles of 0° (per-tendon mean short component T_2_* 0.60 – 0.83 msec) and 55° (per-tendon mean short component T_2_* 0.93 – 1.63 msec) relative to the main magnetic field at 7T[39]. Our values calculated using three TE were of similar magnitude to those reported for short-component T_2_* in the Achilles tendon of healthy volunteers at 7T, which ranged from 0.20 – 0.48 msec[21]. Tendon long axis angle relative to the main magnetic field was not reported in that work, limiting comparison to the patellar tendon due to the different anatomical orientation of the tendons. We did see some non-exponential behavior in the signal decay of the tendon ROI in some cases (as illustrated in Fig 4), which may be due to partial volume averaging with adjacent adipose tissue or blurring where the tendon is too far from isocenter.

The monoexponential T_2_* of cartilage in uninjured and ACL-reconstruction knees, measured with a mix of ultrashort and conventional TE (TE range: 0.032 – 16 msec) at 3T, has been reported to be around 14 – 20 msec[40], and the short/long-T_2_* components of human cadaveric patellar cartilage by biexponential fitting (TE range: 0.08 – 40 msec) at 3T and 1^st^/2^nd^/3^rd^ components of bovine nasal cartilage by triexponential fitting (TE range: 0.6 – 614.4 msec) at 9.4T have been reported to be about 0.50/35 msec and 2.3/25.2/96.3 msec respectively[41]. Our cartilage ultrashort-T_2_* values of 1.0 msec (seven TE) and 0.7 msec (three TE) were measured with a sub-millisecond maximum TE, causing a strong ultrashort component weighting, and as a result are more similar to the reported short-component biexponential and triexponential T_2_*values than the longer monoexponential T_2_* values. Ultrashort-T_2_* components may be more sensitive to some pathology than the long components[23].

Conventional- and ultrashort-T_2_* mapping has been less thoroughly explored in skin than it has been in musculoskeletal tissues, but it is of interest since skin is a high-collagen-content tissue that may be impacted alongside musculoskeletal tissues when collagen structure or function is abnormal. Skin conventional-T_2_*, calculated from images acquired at TE values of 2.8 – 60 msec, has been reported to be 9.8 msec at 1.5T[42,43]. Our measured value of 0.8 msec (seven TE)/0.7 msec (three TE) likely reflects only the shorter components since we only acquired images at sub-millisecond TE values.

Our cortical bone ultrashort-T_2_* median value of 0.7 msec (which was the same when using seven and three TE values) was similar to the previously reported bone specimen ultrashort-T_2_* values of 0.4 – 0.7 msec measured at 7T (TE range: 0.064 – 2.048 msec)[5] and are near the bound-water ultrashort-T_2_* values measured previously at 4.7T[35].

Compared to the tendon, cartilage, skin, and cortical bone, the meniscus and ligament ultrashort-T_2_* had lower coefficients of determination (*R^2^* = 0.77 – 0.82 with seven TE and 0.88 – 0.92 with three TE values). The meniscus ultrashort-T_2_* mapping results for both sets of TE (three and seven TE) are both lower than ultrashort-T_2_* at 3T (7.3 msec)[20] and short-T_2_* values at 7T (7.31 – 9.19 msec)[44] reported previously for healthy menisci. However, our values were within the range of 0.4 – 4 msec reported previously for short-component T_2_* of meniscus *in vitro* at 9.4T[45]. Similarly, our ACL and PCL ultrashort-T_2_* values were lower than the previously reported values[46]. This is likely in part due to the expected reduction in T_2_* at 7T compared to 3T[47]. As noted above, the extent of the field strength-dependence has been shown to vary by region within tendon[21], and this may also be true in the two cruciate ligaments.

The differences between the results here for all tissues and those reported in the literature are also likely due to the difference in TE range used, where most other studies have been performed with either a single or a few ultrashort-TE and a larger number of conventional TE values (>1 msec), in contrast to our use of only ultrashort-TE values. In addition, our use of a coefficient of determination threshold for excluding poor-fit voxels is less commonly used than approaches in which voxels are excluded for having T_2_* or T_2_ values outside of an expected measurement range. In some tissues up to 60% (seven TE fitting) or up to 51% (three TE fitting) of voxels needed to be excluded due to poor fit. While the use of bicomponent exponential fitting and models with incorporation of Gaussian decay and oscillating off-resonance components have been found to improve the fit of tendon[48,49] and meniscus[45] T_2_* signal decay, these more complex models were not explored in this current work, as the upper range of TE values and number of TE points acquired were not sufficient to enable this more complex modeling. However, these components do likely contribute to the signal behavior and signal change between TE values. These factors, along with partial volume averaging and potential issues due to movement between series, artifact, or noise, may contribute to the poor monoexponential fit in some voxels.

### Limitations

This work was limited to imaging of MnCl_2_ and collagen phantoms and a small number of healthy volunteers. Additional image optimization and parameter adjustment may be needed for optimized imaging of patients. The sensitivity and specificity of PETRA-based short-TE signal measurements and ultrashort-T_2_* mapping for quantifying tissue health remains to be assessed. The small sample size and the strong impact on ultrashort-T_2_* of the TE values used here limits the direct comparison between the reported values and our measurements.

PETRA’s combination of pointwise and radial sampling varies depending on TE, and this is expected to have some impact on the measured signal. Future work may be needed to assess the impact on the accuracy of PETRA ultrashort-T_2_* measurements.

Ultrashort-T_2_* mapping using PETRA does require relatively long cumulative scan time and low resolution to avoid unrealistic scan times. We found that the maximum TE needed to be restricted to 0.58 msec to avoid extremely high scan times for single scans. Bicomponent or multicomponent T_2_* mapping for musculoskeletal tissues would require a wider range of TE values, since long-T_2_* tissue suppression by subtraction imaging and quantitative ultrashort-T_2_* mapping require at least one TE long enough, generally >1 msec, for signal of the short-T_2_* tissue of interest to significantly decrease compared to the long-T_2_* tissue values[2,21,41]. A potential solution to the cumulative scan time with long TE values may be to fix the central portions of k-space filled by the single pointwise approach to an upper limit as function of TE. In this manner, scan time would be shortened while maintaining image resolution and SNR.

## Conclusion

In this work we present the results of phantom and in vivo knee imaging using a modified PETRA sequence with an increased TE range. The results obtained are encouraging, demonstrating that following suitable image optimization, this updated PETRA sequence can provide high quality in vivo images. Furthermore, ultrashort-T_2_* mapping was feasible using the modified extended TE range in the phantoms and in vivo in a small sample of healthy knees. Work is in progress to reduce scan time at long TEs by setting the Cartesian central region of the k-space to an upper limit while preserving image information.

## Acknowledgements

We thank Dr. Dean Hoffmeister, MD, for providing input on image interpretation and clinical applications, and the Carle Illinois Advanced Imaging Center 7T MRI technologists who acquired the MR imaging data.

## Supporting information

**S1 Table. PETRA scan parameters selected as optimal for visualizing knee structures within a reasonable scan time.**

## References

1. Breighner RE, Endo Y, Konin GP, Gulotta LV, Koff MF, Potter HG. Technical Developments: Zero Echo Time Imaging of the Shoulder: Enhanced Osseous Detail by Using MR Imaging. Radiology. 2018;286: 960–966. doi:10.1148/radiol.2017170906

2. Chang EY, Du J, Chung CB. UTE imaging in the musculoskeletal system: UTE Imaging in the MSK System. J Magn Reson Imaging. 2015;41: 870–883. doi:10.1002/jmri.24713

3. Froidevaux R, Weiger M, Brunner DO, Dietrich BE, Wilm BJ, Pruessmann KP. Filling the dead-time gap in zero echo time MRI: Principles compared. Magn Reson Med. 2018;79: 2036– 2045. doi:10.1002/mrm.26875

4. Han M, Larson PEZ, Liu J, Krug R. Depiction of Achilles Tendon Microstructure In Vivo Using High-Resolution 3-Dimensional Ultrashort Echo-Time Magnetic Resonance Imaging at 7 T: Invest Radiol. 2014;49: 339–345. doi:10.1097/RLI.0000000000000025

5. Krug R, Larson PEZ, Wang C, Burghardt AJ, Kelley DAC, Link TM, et al. Ultrashort echo time MRI of cortical bone at 7 tesla field strength: a feasibility study. J Magn Reson Imaging JMRI. 2011;34: 691–695. doi:10.1002/jmri.22648

6. Larson PEZ, Han M, Krug R, Jakary A, Nelson SJ, Vigneron DB, et al. Ultrashort echo time and zero echo time MRI at 7T. Magn Reson Mater Phys Biol Med. 2016;29: 359–370. doi:10.1007/s10334-015-0509-0

7. Li C, Magland JF, Zhao X, Seifert AC, Wehrli FW. Selective in vivo bone imaging with long- *T* _2_ suppressed PETRA MRI: Long- *T* _2_ Suppressed PETRA MRI. Magn Reson Med. 2017;77: 989– 997. doi:10.1002/mrm.26178

8. Schieban K, Weiger M, Hennel F, Boss A, Pruessmann KP. ZTE imaging with enhanced flip angle using modulated excitation. Magn Reson Med. 2015;74: 684–693. doi:10.1002/mrm.25464

9. Weiger M, Brunner DO, Dietrich BE, Müller CF, Pruessmann KP. ZTE imaging in humans. Magn Reson Med. 2013;70: 328–332. doi:10.1002/mrm.24816

10. Zibetti MVW, Johnson PM, Sharafi A, Hammernik K, Knoll F, Regatte RR. Rapid mono and biexponential 3D-T1ρ mapping of knee cartilage using variational networks. Sci Rep. 2020;10: 19144. doi:10.1038/s41598-020-76126-x

11. Lee YH, Suh J-S, Grodzki D. Ultrashort echo (UTE) versus pointwise encoding time reduction with radial acquisition (PETRA) sequences at 3 Tesla for knee meniscus: A comparative study. Magn Reson Imaging. 2016;34: 75–80. doi:10.1016/j.mri.2015.09.003

12. Sharafi A, Baboli R, Chang G, Regatte RR. 3D-T _1ρ_ prepared zero echo time-based PETRA sequence for in vivo biexponential relaxation mapping of semisolid short-T _2_ tissues at 3 T. J Magn Reson Imaging. 2019;50: 1207–1218. doi:10.1002/jmri.26664

13. Grodzki DM, Jakob PM, Heismann B. Ultrashort echo time imaging using pointwise encoding time reduction with radial acquisition (PETRA). Magn Reson Med. 2012;67: 510–518. doi:10.1002/mrm.23017

14. Kim SK, Kim D, Lee SJ, Choo HJ, Oh M, Son Y, et al. Clinical value of pointwise encoding time reduction with radial acquisition (PETRA) MR sequence in assessing internal derangement of knee. Clin Imaging. 2018;51: 260–265. doi:10.1016/j.clinimag.2018.05.022

15. Lee HM, Weiger M, Giehr C, Froidevaux R, Brunner DO, Rösler MB, et al. Long-T _2_ -suppressed zero echo time imaging with weighted echo subtraction and gradient error correction. Magn Reson Med. 2020;83: 412–426. doi:10.1002/mrm.27925

16. Weiger M, Brunner DO, Tabbert M, Pavan M, Schmid T, Pruessmann KP. Exploring the bandwidth limits of ZTE imaging: Spatial response, out-of-band signals, and noise propagation. Magn Reson Med. 2015;74: 1236–1247. doi:10.1002/mrm.25509

17. Hager B, Walzer SM, Deligianni X, Bieri O, Berg A, Schreiner MM, et al. Orientation dependence and decay characteristics of T _2_ * relaxation in the human meniscus studied with 7 Tesla MR microscopy and compared to histology. Magn Reson Med. 2019;81: 921–933. doi:10.1002/mrm.27443

18. Williams A, Qian Y, Bear D, Chu CR. Assessing degeneration of human articular cartilage with ultra-short echo time (UTE) T2* mapping. Osteoarthritis Cartilage. 2010;18: 539–546. doi:10.1016/j.joca.2010.02.001

19. Imamura R, Teramoto A, Murahashi Y, Okada Y, Okimura S, Akatsuka Y, et al. Ultra-Short Echo Time–MRI T2* Mapping of Articular Cartilage Layers Is Associated with Histological Early Degeneration. CARTILAGE. 2023; 19476035231205685. doi:10.1177/19476035231205685

20. Williams A, Qian Y, Golla S, Chu CR. UTE-T2∗ mapping detects sub-clinical meniscus injury after anterior cruciate ligament tear. Osteoarthritis Cartilage. 2012;20: 486–494. doi:10.1016/j.joca.2012.01.009

21. Juras V, Zbyn S, Pressl C, Valkovic L, Szomolanyi P, Frollo I, et al. Regional variations of T2* in healthy and pathologic achilles tendon in vivo at 7 Tesla: Preliminary results. Magn Reson Med. 2012;68: 1607–1613. doi:10.1002/mrm.24136

22. Robson MD, Benjamin M, Gishen P, Bydder GM. Magnetic resonance imaging of the Achilles tendon using ultrashort TE (UTE) pulse sequences. Clin Radiol. 2004;59: 727–735. doi:10.1016/j.crad.2003.11.021

23. Wilms LM, Radke KL, Latz D, Thiel TA, Frenken M, Kamp B, et al. UTE-T2* versus conventional T2* mapping to assess posterior cruciate ligament ultrastructure and integrity—an in-situ study. Quant Imaging Med Surg. 2022;12: 4190–4201. doi:10.21037/qims-22-251

24. Jerban S, Hananouchi T, Ma Y, Namiranian B, Dorthe EW, Wong JH, et al. Correlation between the elastic modulus of anterior cruciate ligament (ACL) and quantitative ultrashort echo time (UTE) magnetic resonance imaging. J Orthop Res. 2022; jor.25266. doi:10.1002/jor.25266

25. Keenan KE, Ainslie M, Barker AJ, Boss MA, Cecil KM, Charles C, et al. Quantitative magnetic resonance imaging phantoms: A review and the need for a system phantom: Quantitative MRI Phantoms Review. Magn Reson Med. 2018;79: 48–61. doi:10.1002/mrm.26982

26. Fedorov A, Beichel R, Kalpathy-Cramer J, Finet J, Fillion-Robin J-C, Pujol S, et al. 3D Slicer as an image computing platform for the Quantitative Imaging Network. Magn Reson Imaging. 2012;30: 1323–1341. doi:10.1016/j.mri.2012.05.001

27. NEMA Standards Publication MS 1-2008 (R2014): Determination of Signal-to-Noise Ratio (SNR) in Diagnostic Magnetic Resonance Imaging. National Electrical Manufacturers Association; 2008. Available: https://www.nema.org/docs/default-source/standards-document-library/ms1-2008-r2014-watermarked.pdf?sfvrsn=2101f7b9_2

28. Ilbey S, Jung M, Emir U, Bock M, Özen AC. Characterizing Off-center MRI with ZTE. Z Für Med Phys. 2022; S0939388922000940. doi:10.1016/j.zemedi.2022.09.002

29. Beveridge JE, Machan JT, Walsh EG, Kiapour AM, Karamchedu NP, Chin KE, et al. Magnetic resonance measurements of tissue quantity and quality using T _2_ * relaxometry predict temporal changes in the biomechanical properties of the healing ACL: T _2_ * PREDICTS ACL BIOMECHANICAL PROPERTIES. J Orthop Res. 2018;36: 1701–1709. doi:10.1002/jor.23830

30. Gao J, Xu X, Yu X, Fu Y, Zhang H, Gu S, et al. Quantitatively relating magnetic resonance *T* 1 and *T* 2 to glycosaminoglycan and collagen concentrations mediated by penetrated contrast agents and biomacromolecule-bound water. Regen Biomater. 2023;10: rbad035. doi:10.1093/rb/rbad035

31. Aro O-P. Comparison of relaxation properties of collagen gel MRI phantoms at multiple magnetic fields. University of Oulu. 2020. Available: https://urn.fi/URN:NBN:fi:oulu-202008152795

32. Martin MN, Jordanova KV, Kos AB, Russek SE, Keenan KE, Stupic KF. Relaxation measurements of an MRI system phantom at low magnetic field strengths. Magn Reson Mater Phys Biol Med. 2023;36: 477–485. doi:10.1007/s10334-023-01086-y

33. Stanisz GJ, Odrobina EE, Pun J, Escaravage M, Graham SJ, Bronskill MJ, et al. T _1_, T _2_ relaxation and magnetization transfer in tissue at 3T. Magn Reson Med. 2005;54: 507–512. doi:10.1002/mrm.20605

34. Horch RA, Gore JC, Does MD. Origins of the ultrashort- *T* _2_ ^1^ H NMR signals in myelinated nerve: A direct measure of myelin content?: Ultrashort-T _2_ in Myelinated Nerve. Magn Reson Med. 2011;66: 24–31. doi:10.1002/mrm.22980

35. Ketsiri T, Uppuganti S, Harkins KD, Gochberg DF, Nyman JS, Does MD. T _1_ relaxation of bound and pore water in cortical bone. NMR Biomed. 2023;36: e4878. doi:10.1002/nbm.4878

36. Filho GH, Du J, Pak BC, Statum S, Znamorowski R, Haghighi P, et al. Quantitative Characterization of the Achilles Tendon in Cadaveric Specimens: T1 and T2 * Measurements Using Ultrashort-TE MRI at 3 T. Am J Roentgenol. 2009;192: W117–W124. doi:10.2214/AJR.07.3990

37. Ma Y, Zhao W, Wan L, Guo T, Searleman A, Jang H, et al. Whole knee joint T _1_ values measured in vivo at 3T by combined 3D ultrashort echo time cones actual flip angle and variable flip angle methods. Magn Reson Med. 2019;81: 1634–1644. doi:10.1002/mrm.27510

38. Richard S, Querleux B, Bittoun J, Jolivet O, Idy-Peretti I, De Lacharriere O, et al. Characterization of the Skin In Vivo by High Resolution Magnetic Resonance Imaging: Water Behavior and Age-Related Effects. J Invest Dermatol. 1993;100: 705–709. doi:10.1111/1523-1747.ep12472356

39. Hager B, Schreiner MM, Walzer SM, Hirtler L, Mlynarik V, Berg A, et al. Transverse Relaxation Anisotropy of the Achilles and Patellar Tendon Studied by MR Microscopy. J Magn Reson Imaging. 2022;56: 1091–1103. doi:10.1002/jmri.28095

40. Williams AA, Titchenal MR, Do BH, Guha A, Chu CR. MRI UTE-T2* shows high incidence of cartilage subsurface matrix changes 2 years after ACL reconstruction: MRI UTE-T2* 2 YEARS POST-ACLR. J Orthop Res. 2019;37: 370–377. doi:10.1002/jor.24110

41. Du J, Diaz E, Carl M, Bae W, Chung CB, Bydder GM. Ultrashort echo time imaging with bicomponent analysis. Magn Reson Med. 2012;67: 645–649. doi:10.1002/mrm.23047

42. Song HK, Wehrli FW, Ma J. In vivo MR microscopy of the human skin. Magn Reson Med. 1997;37: 185–191. doi:10.1002/mrm.1910370207

43. Mirrashed F, Sharp JC. *In vivo* morphological characterisation of skin by MRI micro-imaging methods. Skin Res Technol. 2004;10: 149–160. doi:10.1111/j.1600-0846.2004.00071.x

44. Kajabi AW, Zbýň Š, Smith JS, Hedayati E, Knutsen K, Tollefson LV, et al. Seven tesla knee MRI T2*-mapping detects intrasubstance meniscus degeneration in patients with posterior root tears. Radiol Adv. 2024;1: umae005. doi:10.1093/radadv/umae005

45. Kirsch S, Kreinest M, Reisig G, Schwarz MLR, Ströbel P, Schad LR. *In vitro* mapping of ^1^ H ultrashort *T* _2_ * and *T* _2_ of porcine menisci. NMR Biomed. 2013;26: 1167–1175. doi:10.1002/nbm.2931

46. Liu J, Nazaran A, Ma Y, Chen H, Zhu Y, Du J, et al. Single- and Bicomponent Analyses of T2 ⁎ Relaxation in Knee Tendon and Ligament by Using 3D Ultrashort Echo Time Cones (UTE Cones) Magnetic Resonance Imaging. BioMed Res Int. 2019;2019: 1–9. doi:10.1155/2019/8597423

47. Juras V, Schreiner M, Laurent D, Zbýň Š, Mlynarik V, Szomolanyi P, et al. The comparison of the performance of 3 T and 7 T T2 mapping for untreated low-grade cartilage lesions. Magn Reson Imaging. 2019;55: 86–92. doi:10.1016/j.mri.2018.09.021

48. Anjum MAR, Gonzalez FM, Swain A, Leisen J, Hosseini Z, Singer A, et al. Multi-component relaxation modelling in human Achilles tendon: Quantifying chemical shift information in ultra-short echo time imaging. Magn Reson Med. 2021;86: 415–428. doi:10.1002/mrm.28686

49. Liu F, Kijowski R. Assessment of different fitting methods for in-vivo bi-component T2* analysis of human patellar tendon in magnetic resonance imaging. Muscles Ligaments Tendons J. 2017;7: 163–172. doi:10.11138/mltj/2017.7.1.163

